# Comparison of histological delineations of medial temporal lobe cortices by four independent neuroanatomy laboratories

**DOI:** 10.1101/2023.05.24.542054

**Authors:** Anika Wuestefeld, Hannah Baumeister, Jenna N. Adams, Robin de Flores, Carl Hodgetts, Negar Mazloum-Farzaghi, Rosanna K. Olsen, Vyash Puliyadi, Tammy T. Tran, Arnold Bakker, Kelsey L. Canada, Marshall A. Dalton, Ana M. Daugherty, Renaud La Joie, Lei Wang, Madigan Bedard, Esther Buendia, Eunice Chung, Amanda Denning, María del Mar Arroyo-Jiménez, Emilio Artacho-Pérula, David J. Irwin, Ranjit Ittyerah, Edward B. Lee, Sydney Lim, María del Pilar Marcos-Rabal, Maria Mercedes Iñiguez de Onzoño Martin, Monica Munoz Lopez, Carlos de la Rosa Prieto, Theresa Schuck, Winifred Trotman, Alicia Vela, Paul Yushkevich, Katrin Amunts, Jean C. Augustinack, Song-Lin Ding, Ricardo Insausti, Olga Kedo, David Berron, Laura E.M. Wisse

**Affiliations:** Clinical Memory Research Unit, Department of Clinical Sciences, Malmö, Lund University, Sweden; German Center for Neurodegenerative Diseases (DZNE), Magdeburg, Germany; Department of Neurobiology and Behavior, University of California, Irvine, Irvine, CA, USA; INSERM UMR-S U1237, PhIND “Physiopathology and Imaging of Neurological Disorders”, Institut Blood and Brain, Caen-Normandie University, Caen-Normandie, France; Royal Holloway, University of London, Egham, UK; University of Toronto, Toronto, ON, Canada; Rotman Research Institute, North York, ON, Canada; Department of Psychological and Brain Sciences, Johns Hopkins University, Baltimore, MD, USA; Department of Psychology, Stanford University, Stanford, CA, USA; Institute of Gerontology, Wayne State University, Detroit, MI, USA; School of Psychology, University of Sydney, Australia; Department of Psychology, Wayne State University, Detroit, MI, USA; Memory and Aging Center, Department of Neurology, Weill Institute for Neurosciences, University of California, San Francisco USA; The Ohio State University, Columbus, OH, USA; Department of Pharmacology, University of North Carolina at Chapel Hill, Chapel Hill, NC, USA; University of Castilla-La Mancha, Albacete, Spain; University of Pennsylvania, Philadelphia, PA, USA; Institute of Neuroscience and Medicine (INM-1), Research Center Jülich, Jülich, Germany; C. & O. Vogt Institute for Brain Research, Medical Faculty, University Hospital Düsseldorf, Heinrich-Heine-University, Düsseldorf, Germany; Massachusetts General Hospital, Boston, MA, USA; Allen Institute for Brain Science, Seattle, WA, USA; Department of Diagnostic Radiology, Lund University, Sweden

**Author notes:** A.W. and H.B. contributed equally to this work. D.B. and L.E.M.W. contributed equally to this work.

**Keywords:** parahippocampal gyrus, harmonization, neuroimaging, segmentation, entorhinal cortex, Brodmann area 35, Brodmann area 36, parahippocampal cortex

## Abstract

The medial temporal lobe (MTL) cortex, located adjacent to the hippocampus, is crucial for memory and prone to the accumulation of certain neuropathologies such as Alzheimer’s disease neurofibrillary tau tangles. The MTL cortex is composed of several subregions which differ in their functional and cytoarchitectonic features. As neuroanatomical schools rely on different cytoarchitectonic definitions of these subregions, it is unclear to what extent their delineations of MTL cortex subregions overlap. Here, we provide an overview of cytoarchitectonic definitions of the cortices that make up the parahippocampal gyrus (entorhinal and parahippocampal cortices) and the adjacent Brodmann areas (BA) 35 and 36, as provided by four neuroanatomists from different laboratories, aiming to identify the rationale for overlapping and diverging delineations.

Nissl-stained series were acquired from the temporal lobes of three human specimens (two right and one left hemisphere). Slices (50 µm thick) were prepared perpendicular to the long axis of the hippocampus spanning the entire longitudinal extent of the MTL cortex. Four neuroanatomists annotated MTL cortex subregions on digitized (20X resolution) slices with 5 mm spacing. Parcellations, terminology, and border placement were compared among neuroanatomists. Cytoarchitectonic features of each subregion are described in detail.

Qualitative analysis of the annotations showed higher agreement in the definitions of the entorhinal cortex and BA35, while definitions of BA36 and the parahippocampal cortex exhibited less overlap among neuroanatomists. The degree of overlap of cytoarchitectonic definitions was partially reflected in the neuroanatomists’ agreement on the respective delineations. Lower agreement in annotations was observed in transitional zones between structures where seminal cytoarchitectonic features are expressed more gradually.

The results highlight that definitions and parcellations of the MTL cortex differ among neuroanatomical schools and thereby increase understanding of why these differences may arise. This work sets a crucial foundation to further advance anatomically-informed human neuroimaging research on the MTL cortex.

## Introduction

The medial temporal lobe (MTL) cortex – located adjacent to the hippocampus and comprising the ambient and parahippocampal gyri as well as, in some definitions, the fusiform gyrus (Ding & Van Hoesen, 2010; Stenger et al., 2022; Figure 1A) – is not only crucial for memory (Eichenbaum et al., 1994; Ritchey et al., 2015; Squire & Zola-Morgan, 1991) but also for olfaction, attention, spatial navigation, and, social cognition (Gottfried, 2010; Hannula & Duff, 2017; Iizuka et al., 2021; Levy et al., 1997). Clinically, the MTL cortex is a key brain region for neurodegenerative processes, for instance due to Alzheimer’s disease (Berron et al., 2021; Braak & Braak, 1991; Lace et al., 2009; Llamas-Rodríguez et al., 2022; Matej et al., 2019; Yushkevich, Muñoz López, et al., 2021) or limbic-predominant age-related transactive response DNA binding protein-43 encephalopathy (TDP-43; Matej et al., 2019; Nelson et al., 2019; Yushkevich, Muñoz López, et al., 2021). In each of these conditions, early pathological changes occur in the MTL cortex which, in turn, have been related to emerging amnestic symptoms (Braak & Braak, 1991; Nelson et al., 2012, 2019).

**Figure 1.**
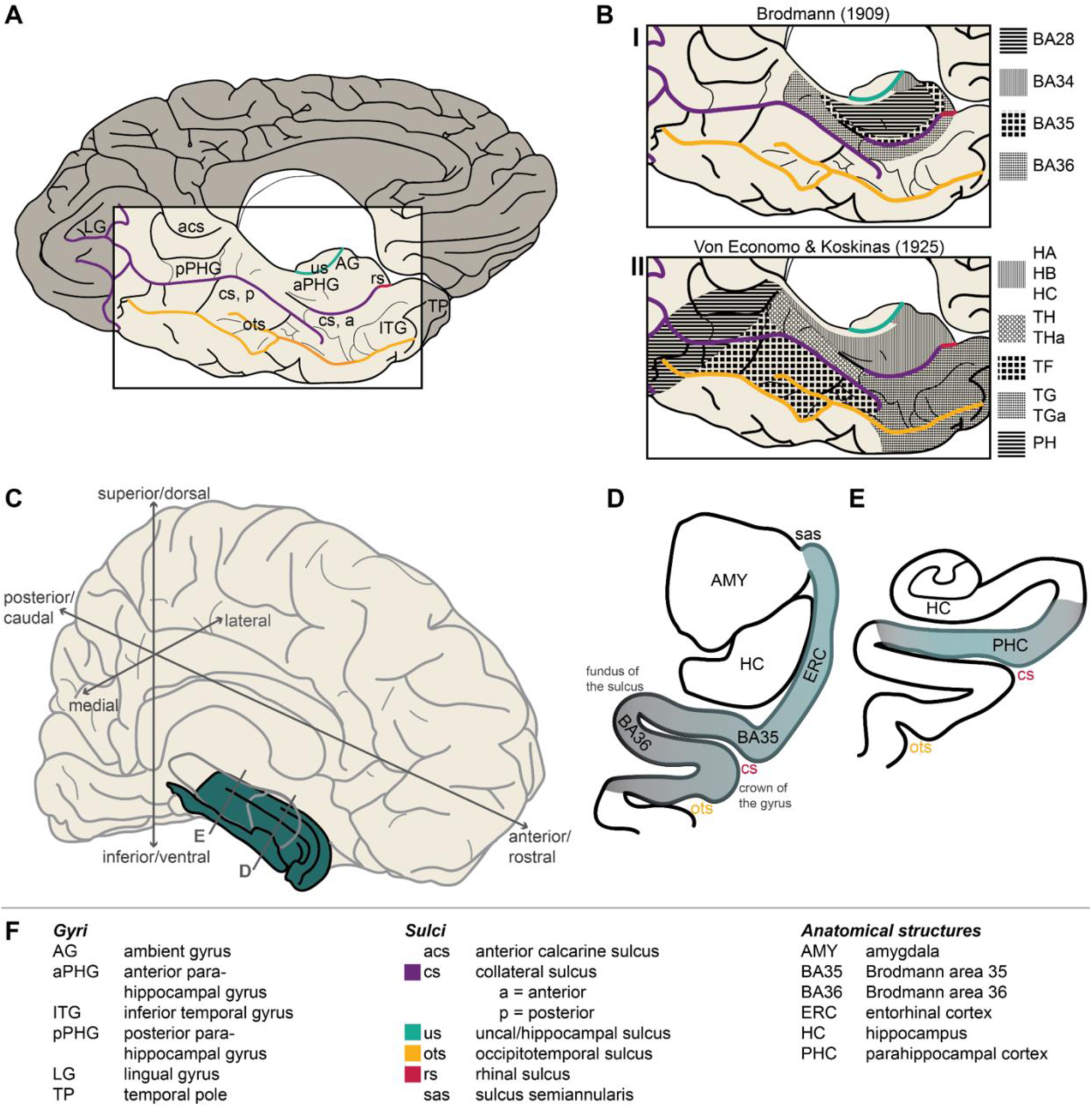
Overview of the cortical medial temporal lobe structures. **A** shows an inferior-medial view on the brain cut at the midline including labels for relevant gyri and sulci. **B** approximates two historical human brain maps (**BI**: Brodmann, 1909; **BII**: von Economo & Koskinas, 1925). **C** is a 3D visualization of the medial temporal lobe cortex (in green) embedded in the left hemisphere indicating important terminology, and identifying the approximate position of subfigures **D** and **E**. **D-E** visualize the rough position of the MTL cortex subregions investigated in this study. **D** shows an anterior coronal section of the MTL cortex including the entorhinal cortex and Brodmann areas 35 and 36 labels. The gradient of color indicates certainty of borders (blue: higher agreement, gray: lower agreement). **E** shows a more posterior section of the MTL cortex showing the parahippocampal cortex. **F** explains the abbreviations used in **A-E**.

Importantly, the MTL cortex is not a single homogeneous entity. It is made up of cortex that forms the gross anatomical parahippocampal gyrus and extends as far as the fusiform gyrus. The cortex of these gyri exhibits high variance in cytoarchitecture. Its anteromedial portion comprises three-layered, allocortical structures of the hippocampus, whereas its outmost lateral subregions exhibit the key characteristics of six-layered isocortex (Braak & Braak, 1995; Ding & Van Hoesen, 2010; Filimonoff, 1947; Insausti et al., 1995, 2017; Van Hoesen et al., 2000). This heterogeneous composition has led neuroanatomists to distinguish various subregions of the human MTL cortex, with partially overlapping terminology.

In their seminal cytoarchitectonic brain maps, Brodmann (1909) as well as von Economo & Koskinas (1925) established two distinct atlases of the MTL cortex (Figure 1B). On Brodmann’s map, it spans Brodmann Area (BA) 28 (*area entorhinalis*), BA34 (*area entorhinalis dorsalis*), BA35 (*area perirhinalis*), and BA36 (*area ectorhinalis*). The term entorhinal cortex (ERC) has been used as an umbrella term for BA28 and BA34, corresponding to its portions located on the anterior parahippocampal and ambient gyri, respectively (Insausti et al., 1995, 2019). The perirhinal cortex is anteriorly, laterally, and posteriorly adjacent to the ERC and corresponds to BA35. BA36 has previously been defined as ectorhinal cortex (Augustinack, Huber, et al., 2013; Brodmann, 1909; Van Hoesen et al., 2000) but has also been included under the term perirhinal cortex (Ding & Van Hoesen, 2010; Insausti et al., 2017; Kivisaari et al., 2012). Importantly, Brodmann’s map is more clear-cut in the anterior MTL cortex but less so in the posterior portion, also referred to as parahippocampal cortex (PHC; not to be confused with parahippocampal gyrus, which is the gross anatomical term for the gyrus adjacent to the hippocampus). In fact, Brodmann (1909) proposed a region termed BA48 (*area retrosubicularis*) to border BA35 posteriorly. However, Brodmann was unable to reliably identify this region in humans. Hence it was not included in Brodmann’s final neuroanatomical atlas, leaving some ambiguity in the posterior MTL cortex. If corresponding to any BA, PHC may include some posterior portions of BA36.

The anatomical map established by von Economo & Koskinas (1925) depicts the MTL cortex as areas HA, HB, HC of the *regio hippocampi* (associated with the entorhinal cortex) and part of TG (TGa) anteriorly (*regio polaris*), as well as regions TH (THa) and TF posteriorly (*regio fusiformis*; Figure 1B). Areas THa and TGa are possible correlates of BA35. THa lies posteromedial to TGa on the crown of the parahippocampal gyrus and adjacent to the parasubiculum or presubiculum of the hippocampal formation. Region TH extends from the lateral crown of the parahippocampal gyrus to the lateral bank of the collateral sulcus and is laterally bordered by the fusiform region TF (von Economo & Koskinas, 1925). Region PH (von Economo & Koskinas, 1929), a large transitional area spanning the posterior parietal and temporal lobes, has been considered part of the PHC as well (Stenger et al., 2022). BA36 may be related to parts of TG posterior TH (*regio temporo-hippocampica*).

Since the establishment of these first exhaustive neuroanatomical atlases, histological studies have benefitted from methodological advancements (e.g., digitalization and immunohistochemistry) and neuroanatomists continuously further developed and specified these existing brain maps. Such advancements led to the distinction of subregions at remarkably detailed levels. For instance, BA35 has been subdivided into BA35a and BA35b based on their distinct cytoarchitecture (Augustinack, Huber, et al., 2013; Ding & Van Hoesen, 2010; Van Hoesen et al., 2000) and several subfields of the ERC have been identified (Behuet et al., 2021; Braak & Braak, 1992; Insausti et al., 1995, 2017; Krimer et al., 1997). A multitude of additional parcellations can be found in the extensive literature on nonhuman primates (e.g., Blatt & Rosene, 1998; Insausti et al., 1987; Suzuki & Amaral, 2003). In addition to the improvement of histological methods, histological datasets have grown. In combination with the rise of *in vivo* neuroimaging techniques, this development has provided crucial insights into interindividual differences in neuroanatomy. For example, it has become apparent that the MTL cortex exhibits great interindividual variation in micro- as well as macrostructure (Augustinack, van der Kouwe, et al., 2013; Ding & Van Hoesen, 2010; Insausti et al., 1998; Ono et al., 1990). In particular, the rhinal and collateral sulci vary in depth and ramification. Variance in collateral sulcus depth and continuity have been linked to differing gross anatomical locations of MTL cortex subregions (Ding & Van Hoesen, 2010; Insausti et al., 1998; Taylor & Probst, 2008). Considering the different histories and subsequent development of anatomical studies, it is unsurprising that neuroanatomical schools rely on different definitions of the MTL cortex.

The described micro- and macroanatomical variance within the MTL cortex poses a challenge to researchers who aim to distinguish MTL cortex subregions on magnetic resonance imaging (MRI) scans for a wide range of research applications. However, accurate delineation is crucial as MTL cortex subregions show distinct involvement in processes of basic and clinical neuroscientific interest. For instance, it has been proposed that the perirhinal cortex and PHC belong to distinct networks of cognition (Maass et al., 2015; Navarro Schröder et al., 2015; Ritchey et al., 2015; Squire & Zola-Morgan, 1991; but also note Doan et al., 2019), making their accurate distinction crucial. With regards to neurodegeneration, subregions of the MTL cortex are differently targeted by pathology and healthy aging (Llamas-Rodríguez et al., 2022; Stark & Stark, 2016). The importance of accurate MTL cortex parcellation becomes evident in the case of Alzheimer’s disease, in which early tau neurofibrillary tangles are located in the transentorhinal region, closely corresponding to BA35 (Braak & Braak, 1985, 1991).

While the neuroanatomical investigation of the MTL cortex is progressing, proposed parcellations and definitions of subregions used for neuroimaging studies continue to diversify (Yushkevich, Amaral, et al., 2015). This diversity may stem from either inter-individual neuroanatomical variability or, as segmentation protocols are informed by different neuroanatomical laboratories, diverging definitions of brain structures. To clarify the influence of the latter, it is necessary to examine the overlap of different neuroanatomical laboratories when delineating brain structures on histological samples. Here, we provide an overview of the cytoarchitectonic definitions of MTL cortex subregions as provided by four neuroanatomists (J.C.A., R.I., O.K., and S.L.D.), based at four different laboratories. We focus on subregions ERC, BA35, BA36, and PHC (Figure 1C-E), as this parcellation is commonly found in the neuroanatomical literature and has been translated into MTL cortex segmentation protocols for MRI (Augustinack, Huber, et al., 2013; Berron et al., 2017; Fischl et al., 2009; Insausti et al., 1998; Xie, Wisse, Pluta, de Flores, et al., 2019; Yushkevich, Pluta, et al., 2015). We additionally compare annotations of these subregions on histological sections of three specimens that all neuroanatomists performed. The insights from this detailed analysis of MTL cortex subregional delineations will provide a basis for the ongoing Hippocampal Subfields Group’s effort to develop a harmonized segmentation protocol for MTL cortex subregions to be used for *in vivo* MRI (Olsen et al., 2019; Wisse et al., 2017; Yushkevich, Amaral, et al., 2015). Moreover, the present study will benefit human neuroimaging studies of the MTL cortex in a broader sense by allowing more precise localization of MTL cortex structures and interpretation of results and ensuring comparability of studies.

## Materials and methods

### Specimens

Three human MTL specimens, two from the Human Neuroanatomy Laboratory (HNL) at University of Castilla-La Mancha (UCLM, Albacete, Spain) and one from the Center for Neurodegenerative Disease Research at the University of Pennsylvania (UPenn, Philadelphia, US), were used. Human brain specimens were obtained in accordance with the UCLM Ethical Clinical Committee, and the University of Pennsylvania Institutional Review Board guidelines. Written consent was obtained in all HNL cases. Where possible, pre-consent during life and, in all cases, next-of-kin consent at death was given at UPenn. The three cases were selected to represent both genders, both hemispheres, a wide age range, cases with and without reported neurodegenerative disease diagnosis, and a broad variability in depth and length of the collateral sulcus to evaluate the location of MTL cortical fields in relationship to macrostructural landmarks. Table 1 provides demographic and diagnostic details of the selected specimens. Note that for ease of interpretation, the left hemisphere specimen (specimen 2) was mirrored in all figures to match the orientation of the other specimens.

**Table 1.** Demographic, diagnostic, and neuropathological information of the three specimens.

|  | Specimen 1 | Specimen 2 | Specimen 3 |
| --- | --- | --- | --- |
| **Age** (years) | 90+<sup>1</sup> | 66 | 83 |
| **Sex** | male | female | female |
| **Postmortem interval**<br>(hours) | 6 | 9 | 4 |
| **Hemisphere annotated** | right | left | right |
| **Collateral sulcus** | deep & continuous | deep & discontinuous | shallow & discontinuous |
| **Diagnosis** | low ADNC | unremarkable brain | low ADNC, PSP |
| **Neuropathological staging** |  |  |  |
| A:B:C score<sup>2</sup> | A1:B1:C0 | A0:B1:C0 | A1:B0:C0 |
| NFT | Braak II | Braak II | Braak 0; 3+ tau (PSP) |

Diagnosis and neuropathological staging were derived from contralateral sampling.<sup>1</sup>As age over 89 is considered protected health information in the US, the exact age of this specimen is not disclosed. <sup>2</sup>Pathology severity rating (A:B:C score; see Hyman et al., 2012). Abbreviations: A=amyloid score based on Thal phases for amyloid deposits; ADNC=Alzheimer's disease neuropathologic change; B=Neurofibrillary tangle stage, modified from Braak & Braak (1991); C=CERAD neuritic plaque score; NFT=neurofibrillary tangles (AT8 staining); PSP=progressive supranuclear palsy.

### Hemisphere preparation and diagnostic pathology

At UCLM, fixation was performed by intracarotid perfusion with the brain in the skull (in situ; Insausti et al., 2023) before brain removal at autopsy with 4% paraformaldehyde in 0.1M phosphate buffer. At UPenn, brain specimens were fixed in 10% neutral buffered formalin after autopsy. For both centers, hemispheres were fixed for at least four weeks. The opposite hemisphere was sampled for diagnostic pathology according to the National Institute of Aging and Alzheimer’s Association (NIA-AA) protocol (Hyman et al., 2012). After hemisphere fixation, an intact tissue block containing the full temporal lobe was dissected and placed in the fixative for 7 tesla (T) MRI scanning.

### Serial histology and immunohistochemistry

Beginning at the temporal pole, specimens were cut coronally following the plane of the 7 tesla MRI into four parallel 20 mm thick blocks using a custom 3D mold that was generated from a postmortem 7 tesla MRI (for details see Yushkevich et al., 2021). Using dry ice, cryoprotected blocks were frozen and serially sectioned into 50 μm sections using a sliding microtome coupled to a freezing unit after digital block-face images were taken.

Every resulting section was collected in a freezing protectant solution. Every 10^th^ section was immediately mounted and Nissl-stained with thionin for cytoarchitectonic analysis, resulting in 0.5 mm gap between adjacent stained sections. The remaining tissue was saved for further use. Every 20^th^ section (1mm gap between stained sections) was stained using AT8, a phosphorylated tau antibody immunohistochemistry stain. All sections were counterstained with lower intensity thionin. Sections were mounted on glass slides, digitally scanned, and uploaded to a cloud-based digital histology archive which supports anatomical labeling (see Supplementary Information).

### Histological annotations

For the purposes of this study, Nissl-stained sections 5 mm apart were selected to be annotated, resulting in a total of 15-16 digitized slices (20X resolution) annotated per specimen. This selection of sections aimed to balance workload for the neuroanatomists while ensuring thorough coverage of the MTL cortex subregions. First, after receiving instruction on the digital annotation tool (see Supplementary Methods), annotations were completed independently by each neuroanatomist (J.C.A., O.K., R.I., S.L.D.). Second, J.C.A., O.K., and R.I. attended a Hippocampal Subfields Group working group meeting in June 2022 during which their annotations and definitions of MTL cortex subregions were discussed. At this meeting every region was discussed individually, comparing the histological annotations of the neuroanatomists – focusing on the main discrepancies and similarities – and asking for clarifications on the reasoning of specific delineations and definitions. Third, J.C.A., O.K., and S.L.D. received access to the full range of serial histology slices to further inform their annotations and potentially adjust them, which two of them completed. R.I. had access to the full range of serial histology slices from the beginning, as part of another project. Fourth, all four neuroanatomists were consulted for follow-up clarifications and asked to provide the seminal cytoarchitectonic features per region that they used for their annotations. Assessments of agreement among neuroanatomists was performed per subregion in a qualitative manner.

## Results

All neuroanatomists determined the borders of ERC, BA35, BA36, and PHC, with finer subdivisions of these subregions made by some neuroanatomists following common practices in their respective laboratories. The following sections first provide a short introduction to the key cytoarchitectonic concepts that informed the neuroanatomists’ annotations. Next, the cytoarchitectonic features, definitions, and parcellations of each MTL cortex subregion, as provided by the neuroanatomists, are described and compared.

### Key cytoarchitectural concepts used for MTL cortex parcellation

Cytoarchitectonic features such as neuronal size, type, organization, and packing density in the different cellular layers were the main source of information used for annotations. At times, some neuroanatomists additionally referred to gross anatomical landmarks in their justifications of border placement. Table 2 provides definitions of key cytoarchitectonic concepts that were relevant for the delineation of MTL cortex subregions.

**Table 2.** Key cytoarchitectonic concepts used for the delineation of the entorhinal cortex, Brodmann areas 35 and 36, and the parahippocampal cortex.

| Concept | Definition | Cortex type | Definition |
| --- | --- | --- | --- |
| Granularity |  | Allocortex | Three-layered cortex |
| Agranular cortex | Lacks granule cell bodies | Periallocortex | Six-layered<sup>1</sup> agranular cortex |
| Dysgranular cortex | Exhibits few loosely packed granule cells | Proisocortex | Six-layered dysgranular cortex in which not all isocortical features are clearly expressed |
| Granular cortex | Prominent and densely packed layer of granule cells | Isocortex | Six-layered granular cortex showing an organized internal and external granule cell layer |
| *Lamina dissecans* | Thoroughly agranular layer splitting the cortex in two halves |  |  |
| Layer II cell islands | Darkly stained stellate cells occurring in clusters in ERC |  |  |
| Columnarity | Columnar organization of cortical cells perpendicular to the cortical layers |  |  |
| Internal vs. external layers | Deep layers close to white matter vs. superficial layers close to pial surface |  |  |

<sup>1</sup>Note that terminology describing periallocortical cell layers may differ from terminology used for (pro-)isocortical regions (see for example Ding & Van Hoesen, 2010), while this paper uses consistent terminology. Additionally, note that the (peri-)allocortical layers are not homologous to the layers of the isocortex, since the (peri-)allocortical layers are evolutionarily different from those of the isocortex. Abbreviations: ERC=entorhinal cortex.

An important concept was granularity, which here focuses on the number and organization of granule cells in layer IV (or the equivalent in ERC; see sTable 1) of the cortex (Figure 2). The degree of granularity can range from the complete lack of granule cell bodies (agranular cortex) to the presence of few, loosely arranged granule cells (dysgranular cortex), to the dense packing of highly organized granule cells (granular cortex; Brodmann, 1909; Insausti et al., 2017; von Economo & Koskinas, 1925). Changes in granularity are one of the main distinguishing features of MTL cortex subregions which reflect the transition from periallocortex (agranular), proisocortex (dysgranular), to isocortex (granular; Brodmann, 1909; Ding & Van Hoesen, 2010; Insausti et al., 2017; von Economo & Koskinas, 1925). A cytoarchitectonic feature that can coincide with a lacking granule cell layer IV is the *lamina dissecans*. It is a thoroughly cell-sparse, agranular layer and, in the MTL cortex, specific to ERC. It divides the cortex into two halves or, if two *laminae dissecans* are present, enclosing a small layer of large pyramidal cells between two cell-free layers (layers Va and Vb in Insausti et al., 1995; see also Insausti et al., 2017; Stephan, 1975). Another concept used to parcellate MTL cortex subregions was the presence or absence of layer II cell islands (Figure 2A). These prominent clusters of large stellate cells, also termed pre-α cell islands (Braak & Braak, 1985), are most prominent in the ERC (Ding & Van Hoesen, 2010; Insausti et al., 1995; Schön et al., 2022). Lastly, columnarity is a feature that is differently expressed throughout layer II of the MTL cortex subregions (Figure 2B). It can be defined as the irregular arrangement of cells forming cell accumulations that are oriented perpendicular to the pial surface and are separated by cell-sparse parts.

**Figure 2.**
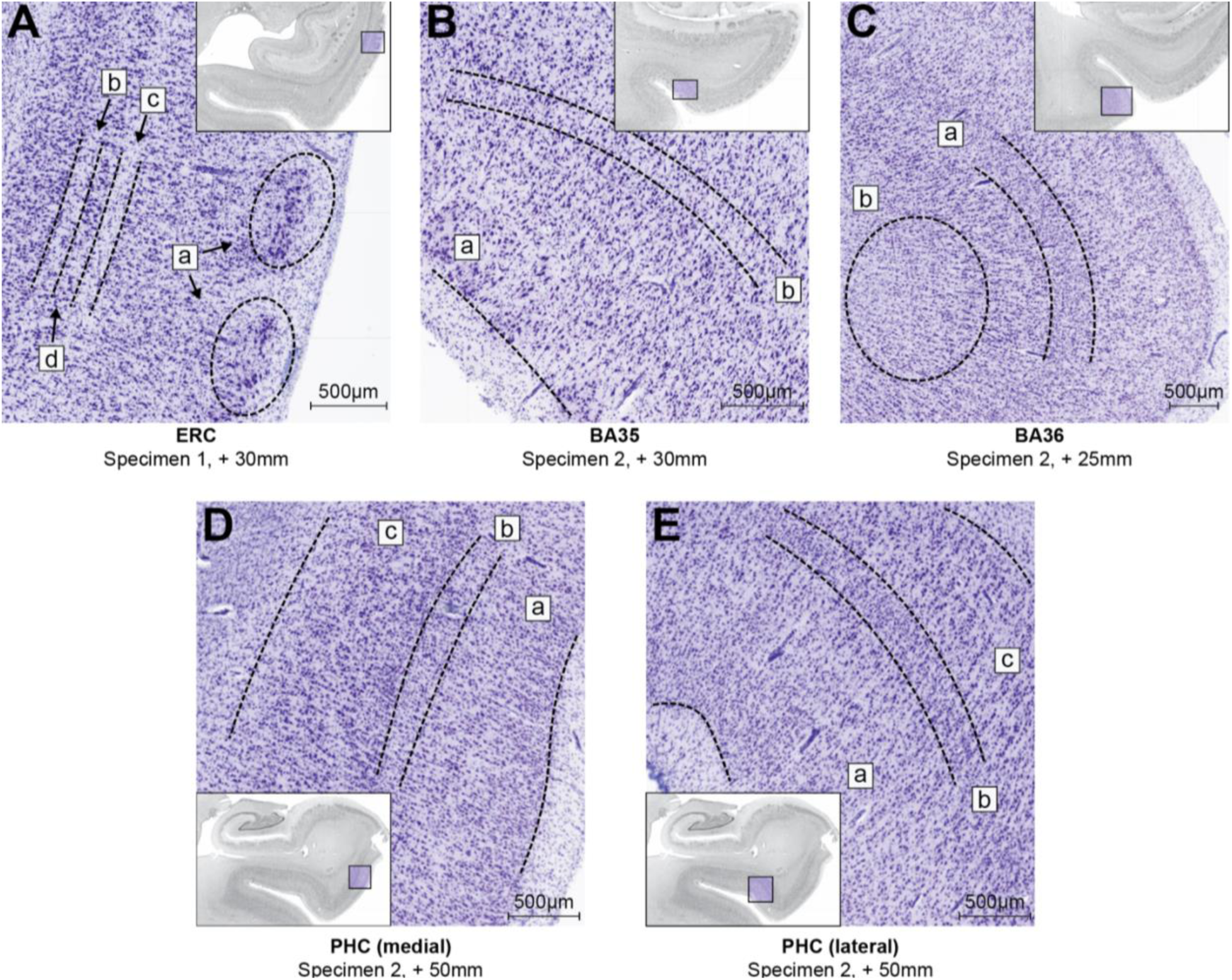
Examples for seminal features for each of the annotated MTL cortex subregions. Millimeters indicate distance from the temporal pole. All displayed examples were annotated as the indicated subregions by all neuroanatomists. **A** shows a section of ERC. Seminal features include layer II cell islands (a) and an internal (b) and external (c) lamina dissecans, which enclose a dense layer of pyramidal cells (d; see sTable 1 for differences in nomenclature). **B** shows a section that was annotated BA35 and shows a columnar organization of layers II and III (a) and an incipient dysgranular layer IV (b). **C** displays a section annotated BA36. It features the region’s thick layer IV, but with loosely organized cells (a) and a fluent transition of layer VI with the white matter (b). PHC was described as a highly heterogeneous region, thus two examples are shown. A more medial section of PHC is shown in **D**, with layer II gradually transitioning into layer III (a), an incipient dysgranular layer IV (b), in which scarce granular cells are intermingled with cells from the neighboring layers, and homogenous layers V and VI (c). Features in more lateral parts of the PHC, as shown in **E**, are similar. Note that in contrast with the medial section, layer II and layer III are more distinguishable (a) and layer IV exhibits an almost isocortex-like degree of granularity (b). Still, no clear border between layers V and VI is visible (c). Note that individual annotations often extended further lateral than shown in **E**. However, there was no slice in which more lateral sections were annotated PHC by all neuroanatomists. Abbreviations: BA=Brodmann area; ERC=entorhinal cortex; PHC=parahippocampal cortex.

### Cytoarchitectonic and gross anatomy features of MTL cortex subregions

Annotations of all neuroanatomists primarily relied on cytoarchitectonics and not on the gross anatomical appearance of the MTL cortex. Only one neuroanatomist additionally referred to gross anatomical hallmarks as a relevant source of information for border definition, as described previously (Ding & Van Hoesen, 2010). Table 3 provides a short overview of each structure’s core cytoarchitectonic features as stated by the neuroanatomists. A more detailed overview can be found in Table 4, which also highlights points of agreement and disagreement among the neuroanatomists. Although the neuroanatomists agreed upon the presence of these features, they were weighted differently by the neuroanatomists when making boundary placement decisions.

**Table 3.** Key features of the MTL cortex subregions.

|  | ERC | Perirhinal | Ectorhinal | PHC |
| --- | --- | --- | --- | --- |
| **Brodmann area** | 28, 34 | 35 | 36 | - |
| **Cortex type<sup>1</sup>** | Periallocortex | Periallocortex and/or proisocortex | Proisocortex or isocortex | Periallocortex, proisocortex, and isocortex |
| **Location<sup>2</sup>** | Crown of the parahippocampal and ambient gyri | Medial and/or lateral bank of the CS | CS and/or crown of the fusiform gyrus | Crown of the parahippocampal gyrus, medial bank of the CS |
| **Seminal feature** (see Figure 2) | - layer II cell islands<br>- *lamina dissecans*<sup>4</sup><br>- layer of large pyramidal cells adjacent to one or two *laminae dissecans*<sup>4</sup> | - columnar organization of layers II and III<br>- no *lamina dissecans*, but agranular, and/or dysgranular layer IV | - increasing granularity in layer IV<br>- gradual transition of layer VI to white matter | - wide internal layers<br>- agranular (medial) to granular (lateral) layer IV<br>- gradual transition between layers V and VI |
| **Agreement<sup>5</sup>** | High | Intermediate to high | Intermediate | Low to intermediate |

<sup>1</sup>For the perirhinal and ectorhinal cortices there was disagreement on the type of cortex (indicated by “or”). <sup>2</sup>The gross anatomical location of subregions varies and is, among other things, dependent on the depth and ramification of the collateral sulcus (see Ding & Van Hoesen, 2010). <sup>3</sup>See Figure 2 for details. <sup>4</sup>Note that some neuroanatomists only refer to the external cell-free layer as *lamina dissecans*. <sup>5</sup>See Table 4 for details. Abbreviations: BA=Brodmann area; CS=collateral sulcus; ERC=entorhinal cortex; PHC=parahippocampal cortex; MTL cortex=parahippocampal gyrus.

**Table 4.** Agreement and disagreement among neuroanatomists about cytoarchitectonic features of the MTL cortex subregions.

| Region | Agreement | Disagreement |  |  |  |
| --- | --- | --- | --- | --- | --- |
|  |  | J.C.A. | O.K. | R.I. | S.L.D. |
| ERC | **Periallocortex**<br>L II<br>Clusters (or islands) of large stellate cells, interspersed with cell free areas<br>L III<br>Wide layer with columnar organization of pyramidal cells<br>L III & IV<br>*Lamina dissecans* formed by substrata of L III & IV (not both are equally present along the longitudinal extent of ERC)<br>L V<br>Big, darkly stained pyramidal neurons<br>L V & VI<br>No clear separation of layers (also with white matter) at anterior levels, clear boundary at posterior levels |  |  |  |  |
| BA35 | Wedge-like transition of lateral ERC to medial BA35 that extends from superficial cell layer to midst of the cortex<br>L II & III<br>Relatively large cells, although not as large as in ERC<br>Columnar organization<br>L III<br>Presence of an oblique band of dark pyramidal cells<sup>1</sup><br>L V<br>Large pyramidal cells | **Periallocortex and proisocortex**<br>L IV<br>Agranular (medial) to dysgranular (lateral) sections | **Periallocortex**<br>L IV<br>Agranular | **Proisocortex**<br>L IV<br>Agranular to dysgranular | **Periallocortex and proisocortex**<br>L IV<br>Agranular (medial) to dysgranular (lateral) sections |
| BA36 | L II<br>Small cells relative to ERC and BA35<br>L III-V<br>Higher cell density in lateral vs. medial portions<br>L III<br>Small pyramidal cells relative to ERC and BA35<br>L V<br>Homogeneous with middle-sized to large pyramidal cells<br>L VI<br>Fluent transition with white matter | **Isocortex**<br>L IV<br>Granular | **Proisocortex**<br>L III<br>Sublamination of L III into L IIIa-c (clearer laterally)<br>L IV<br>Dysgranular<br>L V<br>Higher cell density in the lateral extensions | **Proisocortex**<br>L IV<br>Dysgranular | **Isocortex**<br>L IV<br>Granular |
| PHC | **Periallocortex transitioning to proisocortex**<br>*TH/medial PHC<sup>2</sup>*<br>**Periallocortex**<br>L II<br>- Can be patchy<br>L II & III<br>- Soft transition, superficial cells in L III are similar to LII<br>L IV | Annotated: *TH & TF*<br>*TF*<br>L III<br>Homogeneous<br>Soft transition to L IV | Annotated: *PHC, Ph1-3*<br>*lateral PHC*<br>L III<br>Sublamination of L III into L IIIa-c<br>L IIIa contains small and densely packed cells, L IIIb has least cell density and | Annotated: *TH & TF*<br>*TF*<br>L III<br>Less columnarity than isocortex<br>L IV<br>Thickens caudally | Annotated: *TH & TL/TFm*<br>*TL/TFm (medial)*<br>L IV<br>Soft transition to L V |
|  | - Agranular<br>L V<br>- Thinner compared to TF<br>- Highly developed with large cells<br>L V-VI<br>- Soft transition but smaller and more densely packed cells in L VI<br>- Higher cell density than BA35<br>*TF/TL/TFm/lateral PHC<sup>3</sup>*<br>**Proisocortex**<br>L II<br>- Thicker than TH/medial PHC<br>L III<br>- Higher cell density than BA36<br>L IV<br>- Dysgranular<br>L V<br>- Highly developed with large cells<br>L V-VI<br>- Soft transition but more pronounced than in medial PHC |  | relatively small cells, L IIIc contains middle-sized cells<br>L IV<br>Soft transition to L V<br>*Ph1-3*<br>**Isocortex**<br>For details see Stenger et al. (2022) |  |  |

<sup>1</sup>Note that some but not all neuroanatomists refer to this layer as IIIu. <sup>2</sup>Medial PHC was used by O.K. and TH by the other neuroanatomists. <sup>3</sup>TF can be divided into a medial and lateral part (TFm and TFI). TL is synonymous with TFm. TF was annotated by J.C.A. and R.I. TL/TFm was annotated by S.L.D. O.K. referred to this region as lateral PHC. Abbreviations: BA=Brodmann area; ERC=entorhinal cortex; L=layer; PHC=parahippocampal cortex.

### Entorhinal cortex

Among all subregions, the cytoarchitectonic definition of the ERC exhibited the highest agreement among neuroanatomists. ERC was consistently referred to as a six-layered, periallocortical structure (Augustinack, Huber, et al., 2013; Ding & Van Hoesen, 2010; Insausti et al., 1995, 1998). Compared with the adjacent BA35, its cells are overall relatively large. Layer II comprises large stellate cells (recently also referred to as fan cells; Nilssen et al., 2018; Vandrey et al., 2020) that cluster to form cell islands (Braak & Braak, 1985; Rose, 1927; Figure 2A). These layer II cell islands are highly characteristic for ERC, though they are most prominently expressed in its posteriolateral extent (Behuet et al., 2021). Layer III of ERC is rather thick with pyramidal cells clustered into columns. Neuroanatomists agreed that ERC is a thoroughly agranular structure with a clearly visible *lamina dissecans*, splitting the cortex into an internal (or deep, between white matter and *lamina dissecans*) and external (or superficial, between pial surface and *lamina dissecans*) principal layer (Braak & Braak, 1985; Insausti et al., 2017; Rose, 1927). In some sections of the ERC, two cell-free layers are formed: an external (sometimes referred to as proper *lamina dissecans*) and an internal cell-free layer (sometimes also referred to as *lamina dissecans*, while not by all neuroanatomists; see sTable 1). Note that the terminology used to distinguish cellular layers in the ERC differs from that used in most of the cortex due to ontogenetic differences (Behuet et al., 2021; Braak, 1972). This special case of the ERC has been resolved differently by neuroanatomical laboratories with some staying closer to conventional cortical nomenclature than others (see sTable 1 for an overview; Behuet et al., 2021; Braak & Braak, 1991; Insausti et al., 1995). Between the cell-free layers lies a distinct layer of large, darkly stained pyramidal neurons (Braak & Braak, 1985; Insausti et al., 2017; Rose, 1927). Layers V and VI are close together and do not show sharp boundaries between each other and with the parahippocampal white matter (angular bundle) at anterior and intermediate levels, while posterior ERC subfields show a sharp boundary with the white matter of the angular bundle (Insausti et al., 2017).

Annotations of the ERC along its longitudinal axis are exemplified in Figure 3A-C and their overlap in specimen 1 is visualized in Figure 4A. Generally, there was high agreement on border placement particularly in the midsection of the ERC with slight variability shown at its anterior and posterior ends (Figure 3A and C, Figure 4A). Indeed, neuroanatomists stated that the seminal cytoarchitectonic features of the ERC emerge only gradually at these anterior and posterior transitional sections, making annotations more difficult. An example of gradually developing layer II islands and *lamina dissecans* in the anterior ERC is shown in Figure 5A. In line with the previous neuroanatomical literature, the ERC was consistently bordered anteriorly by the rhinal/collateral sulci and sulcus semiannularis as well as superiorly by the periamygdaloid cortex, reflecting the substantial cytoarchitectonic differences between six-layered ERC the adjacent three-layered periamygdaloid cortex (Insausti et al., 2017). The medial border of ERC was consistently placed adjacent to the hippocampal fissure, as well as the pre- or parasubiculum (Figure 3B-C, Figure 4A; see also Ding & Van Hoesen, 2010; Insausti et al., 1995, 1998, 2017). As already depicted by Brodmann (1909), ERC was encapsulated by perirhinal BA35 at its anterior, posterior, and lateral ends in all three specimens. Compared with the highly consistent superior and medial borders, there was some, albeit minor, variability in the placement of the ERC’s lateral border with BA35, potentially due to the gradual transition of cytoarchitectonic features between ERC and BA35 in this region.

**Figure 3.**
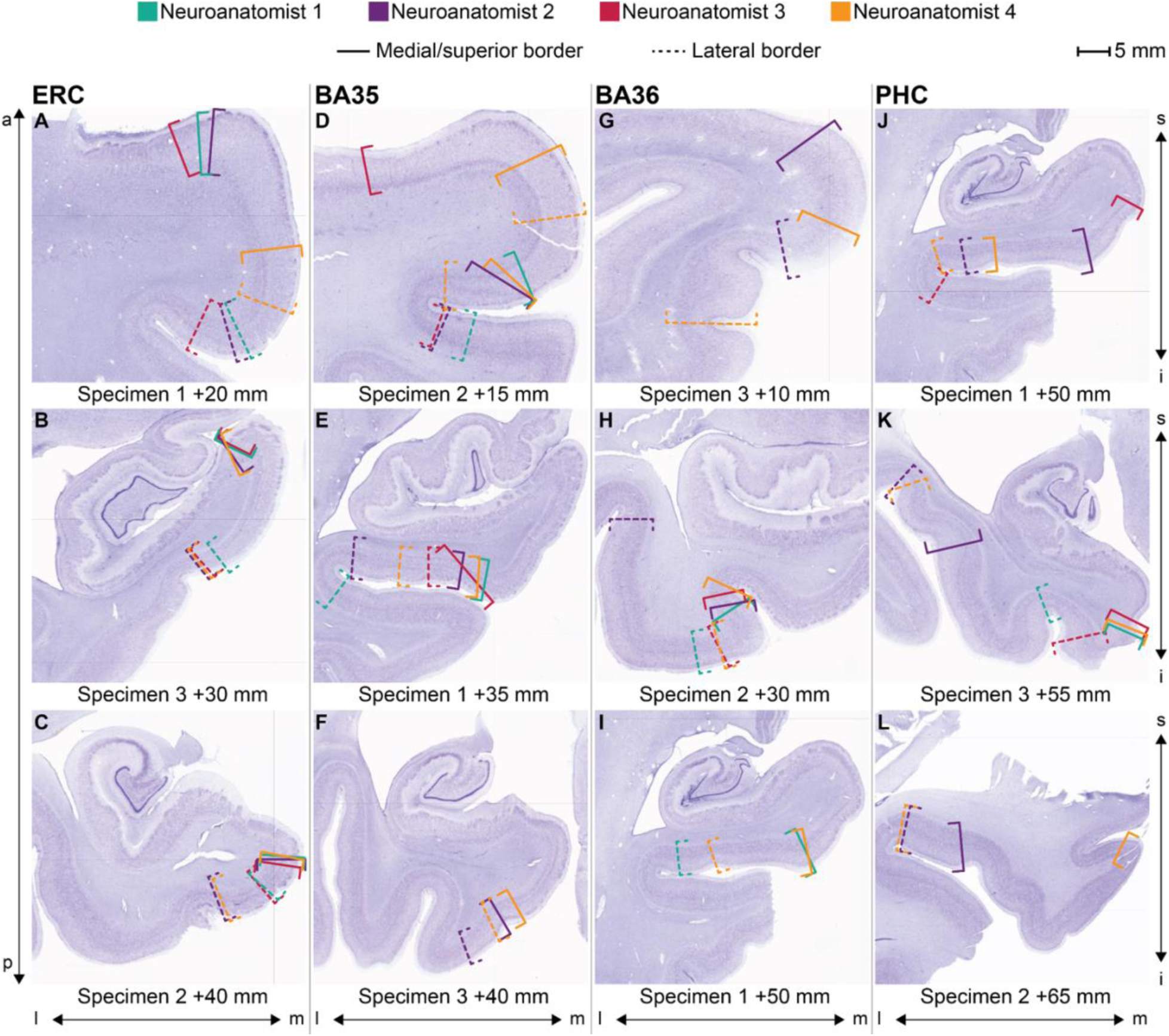
Annotations performed by the neuroanatomists for the anterior, mid-section, and posterior parts of four subregions of the MTL cortex. Millimeters indicate distance from the temporal pole. Sections were chosen to reflect the varying overlap of annotations across specimens and gross anatomical locations within the MTL cortex. Abbreviations: a=anterior; BA=Brodmann area; ERC=entorhinal cortex; i=inferior; l=lateral; m=medial; p=posterior; PHC=parahippocampal cortex; s=superior.

**Figure 4.**
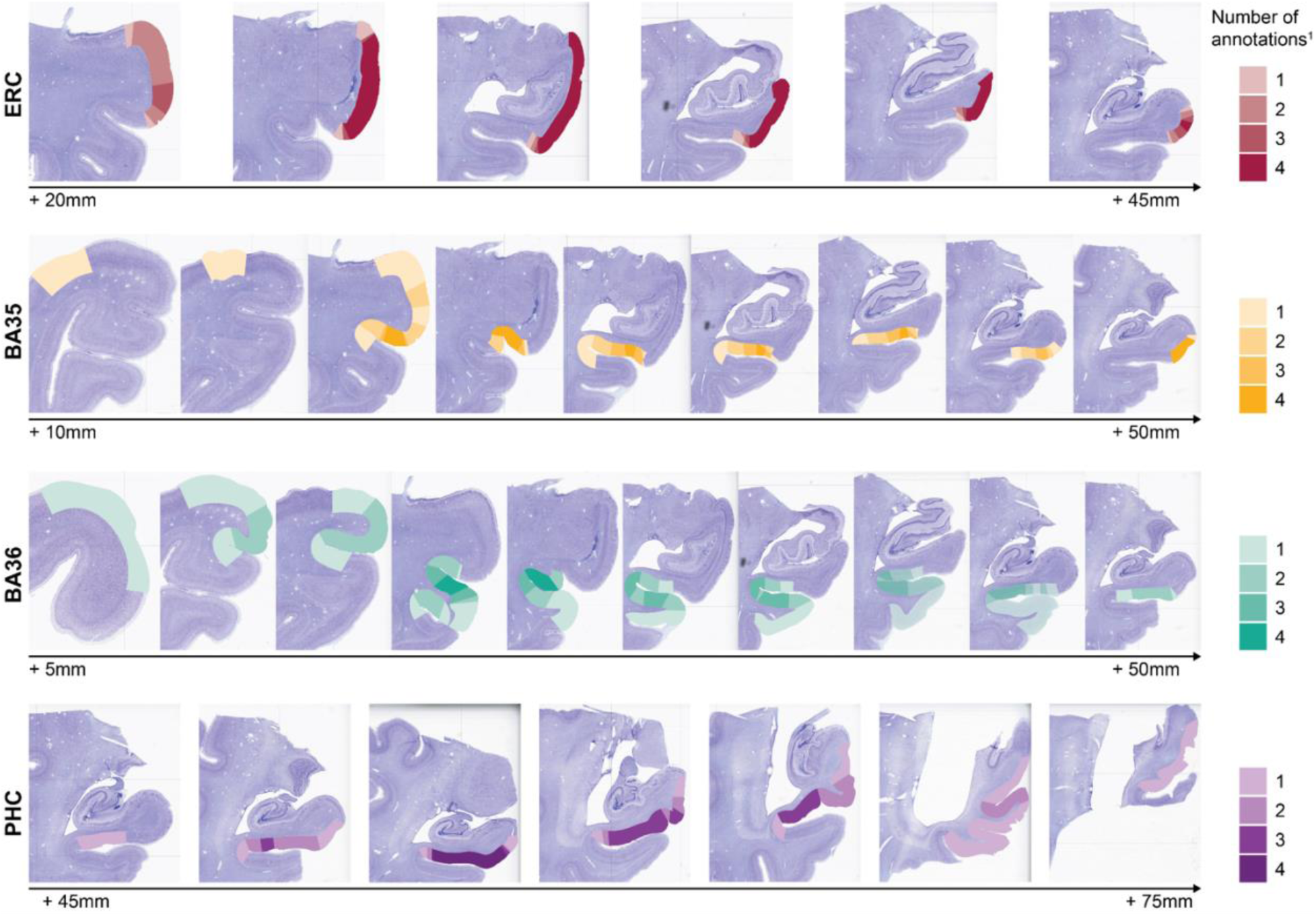
Overview of annotations of all four MTL cortex subregions in Specimen 1 visualizing the overlap among neuroanatomists. Note the lower disagreement (i.e., lighter colors) on transitional slices compared with higher agreement (i.e., darker colors) in midsections. ^1^Indicates how many neuroanatomists annotated a certain part of the cortex as belonging to the specific MTL cortex subregion. Millimeters indicate distance from the temporal pole. Abbreviations: BA=Brodmann area; ERC=entorhinal cortex, PHC=parahippocampal cortex.

**Figure 5.**
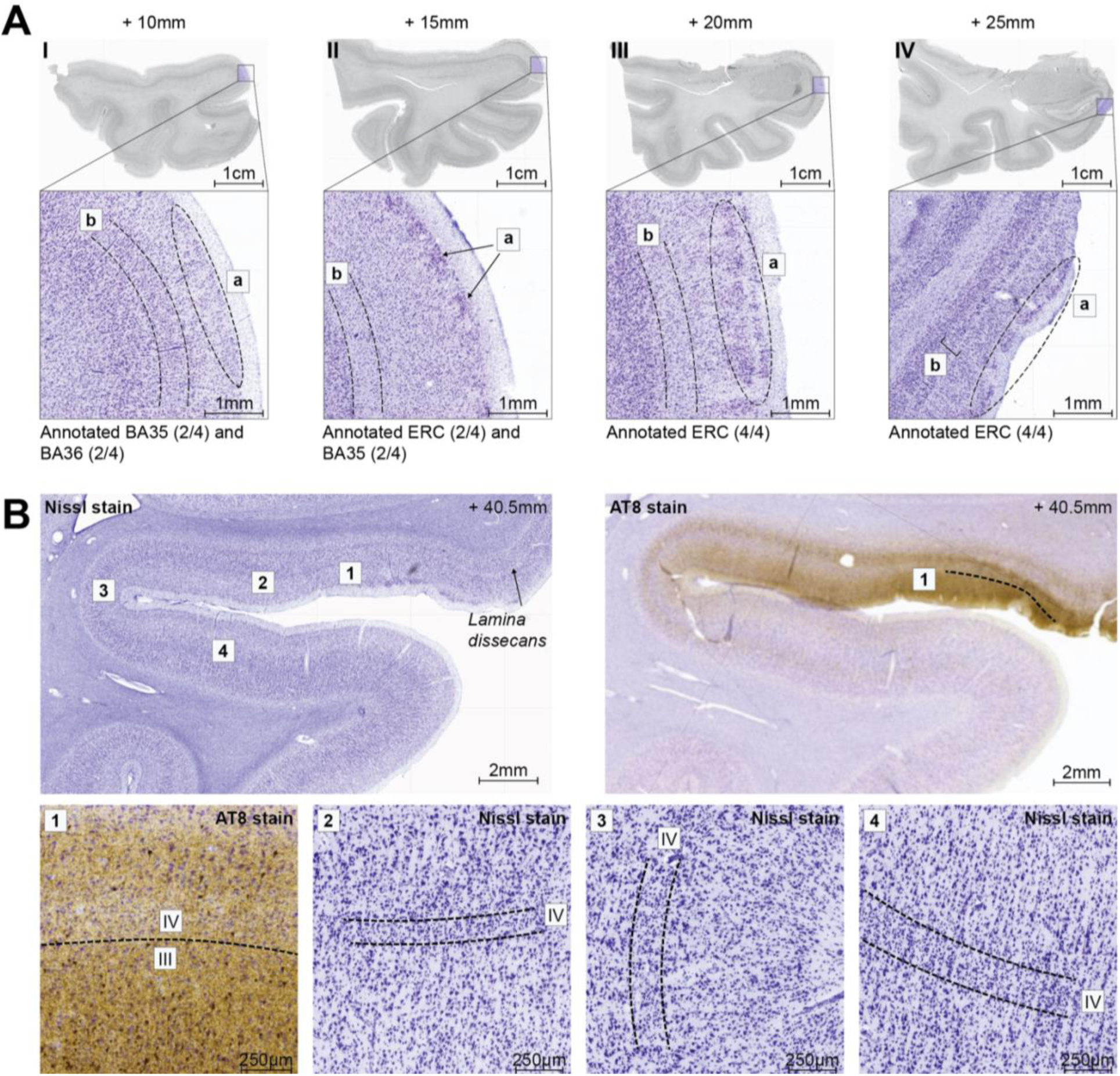
Gradual transition of cytoarchitectonic features in the MTL cortex. **A** shows the gradual transition of the seminal cytoarchitectonic features of ERC in an anterior-to-posterior direction in the anterior MTL cortex in specimen 2 (**A-I** to **A-IV**). Layer II cell islands are increasingly expressed moving posteriorly (a in **A-I** to **A-IV**). At the anterior level (b in **A-I**), layer IV comprises granule cells. In **A-II** and **A-III** two cell-free sublaminae (*laminae dissecans*) can be identified. Two *laminae dissecans* are visible in the very posterior slice (b in **A-IV**). Note that some neuroanatomists only refer to the external cell-free sublamina as *lamina dissecans*. The annotations of the respective section are indicated below each slice. **B** shows the gradual change from *lamina dissecans* (periallocortex) to a fully expressed granular layer IV in the medial-to-lateral direction along the collateral sulcus (cs) in specimen 1. AT8 staining in **B** (right panel) exemplifies the wedge-like appearance of layer III (medial dotted line), made up of uniquely large pyramidal cells (by some neuroanatomists referred to as layer IIIu), which are characteristic for BA35, with the use of AT8 tau staining in the same case. **B1** shows a magnified image of the unique layer III in medial BA35 using AT8 staining. **B2-4** indicate the increasing thickness, cell density, and organization of layer IV moving laterally along the collateral sulcus. Neuroanatomists annotated the section shown in **B2** BA35 (2/4) and BA36 (2/4), in **B3** BA36 (3/4) and transition from BA35 to BA36 (1/4), and in **B4** BA36 (3/4) and non-MTL cortex (1/4). Note: annotations were made on the neighboring slice 0.5 mm anterior to the displayed section. Millimeters indicate distance from the temporal pole. Abbreviations: BA=Brodmann area; ERC=entorhinal cortex.

### Brodmann Area 35

All neuroanatomists agreed that BA35, also called perirhinal cortex, is a heterogeneous structure located at the transition of periallocortex (i.e., ERC) to proisocortex. However, there was disagreement about the specific cortex type of BA35 as it was labeled dysgranular proisocortex by some, while others described both agranular, periallocortical (e.g., BA35a) and dysgranular, proisocortical (e.g., BA35b) portions within BA35 (see Table 3, Table 4). Nevertheless, neuroanatomists agreed on several cytoarchitectonic features being characteristic to BA35. The pyramidal cells found in layers II and V are relatively large, while not as large as in ERC. Layers II and III are organized in a distinct columnar fashion (Figure 2B). BA35 shows a unique layer III (by some neuroanatomists referred to as layer IIIu; Ding et al., 2009) which is formed by exceptionally large cells and lies internally to a typical layer III (see Figure 5B-2; Augustinack, Huber, et al., 2013; Ding et al., 2009; Ding & Van Hoesen, 2010). At the medial border with ERC, these features are expressed to form an oblique border, first appearing superficially, and extending until the midst of the cortex in a lateral direction. Layer IIIu is difficult to distinguish from typical layer III using Nissl-stain, especially for non-experts (Figure 5B; Ding & Van Hoesen, 2010). However, as the pyramidal cells in layer IIIu are affected early by Alzheimer’s disease-related neurofibrillary tangles (Braak & Braak, 1985; Ding & Van Hoesen, 2010), tau staining impressively revealed the unique, wedge-like shape of BA35’s medial border in specimen 2 (Figure 5B-1). BA35 closely corresponds to Braak’s transentorhinal region (Braak & Braak, 1985). While for some neuroanatomists these terms are synonymous and refer to the same region, others stated that only the medial portions of their BA35 annotations (e.g., BA35a) can be referred to as transentorhinal cortex.

Across annotations, BA35 borders BA36 anterolaterally and PHC posterolaterally. Like ERC, annotations in midsections of BA35 exhibited higher agreement among neuroanatomists (Figure 3E-F, Figure 4), compared with anterior (Figure 3D) or posterior transitional sections. Figure 3 and Figure 4 also show that, compared with annotations of the ERC, there was more variation among annotations in the medial-lateral dimension. Specifically, the lateral border of BA35 with BA36 varied among neuroanatomists. This was likely driven by the neuroanatomists’ differing definitions of BA35 with regards to layer IV granularity and cortex type (Table 4) and, importantly, by the gradual expression of layer IV granule cells when moving laterally (Figure 5B). Meanwhile, the medial border with ERC showed high agreement with minor variability (Figure 3D, Figure 4), representing the large consensus about the cytoarchitectonic features of BA35 and ERC in this region (i.e., cell islands in layer II of ERC *versus* large cells in layer II/III arranged in columns in BA35).

### Brodmann Area 36

BA36 was also referred to as ectorhinal cortex by the neuroanatomists. Like BA35, neuroanatomists disagreed on the cortex type and thereby the extent of the layer IV granularity for BA36. While two neuroanatomists denoted BA36 as dysgranular proisocortex, others considered it a fully granular, isocortical structure. The neuroanatomists agreed that layers II and III have smaller cells relative to ERC and BA35, layer V is homogenous with middle to large pyramidal cells, and layer VI fluently transitions into white matter (Figure 2C).

BA36 borders other regions of the temporopolar cortex anteriorly and PHC as well as the peristriate cortex**/**BA19 posteriorly. Higher disagreement among neuroanatomists could be observed in these anterior and posterior transitional regions as opposed to the midsection of BA36 (Figure 3G-I, Figure 4). Among all described structures, the annotations of BA36 exhibited the highest degree of disagreement in the medial-lateral dimension. The lateral border with BA20 varied profoundly. While three neuroanatomists included the lateral bank of the collateral sulcus (i.e., corresponding to Bordmann’s original mapping of BA36), the annotations of one neuroanatomist additionally spanned the entire fusiform gyrus and medial bank of the occipitotemporal sulcus (i.e., also including some of BA20 on Brodmann’s map or area TF as defined by von Economo and Koskinas; Figure 3H, Figure 4). Similar to the border of BA35 with BA36, disagreements on the lateral border of BA36 with BA20 were at least partially caused by the gradual changes in granularity, which is susceptible to individual interpretation, along the medial-lateral axis (Figure 5B) and the neuroanatomists’ different definitions of BA36 as either a proisocortical or isocortical structure (Table 4).

### Parahippocampal Cortex

The PHC is another highly heterogeneous MTL subregion that spans periallocortical and proisocortical portions. In fact, one neuroanatomist stated that PHC even includes granular isocortex at its lateral end. Importantly, the neuroanatomists agreed that parcellation based on granularity is more difficult in the PHC than in more anterior subregions of the MTL cortex. Analogous to more anterior parts of the MTL cortex comprising ERC and BA35, the medial bank of the collateral sulcus exhibits higher cytoarchitectonic heterogeneity at the level of the PHC. Neuroanatomists used various cytoarchitectonic definitions of the PHC which led to substantial differences in annotations, especially in their posterolateral extent (Table 4). In line with many of the previously proposed parcellations of the PHC in nonhuman primates (Blatt & Rosene, 1998; Ding & Van Hoesen, 2010; Suzuki & Amaral, 2003; Tranel et al., 1988) and in humans (Bailey & von Bonin, 1951; von Economo & Koskinas, 1925), its medial portion was referred to as region TH by two of the neuroanatomists. Here, layer II and III cell bodies are rather small, layer IV is absent (agranular), and layers V and VI are homogeneously transitioning into one another (Figure 2D; note that here an incipient granule cell layer may be visible). Relative to BA35, the medial PHC has a higher cell density and wider internal layers (Figure 2D). Neuroanatomists who distinguished PHC subregions divided the lateral PHC into either as TFm (medial) and TFl (lateral; as introduced in the Macaca fascicularis monkey by Suzuki & Amaral, 2003) or as TL (as described also in primates by Blatt & Rosene, 1998). Here, layer II is thicker than in the medial PHC, layer III exhibits high columnarity, weakly organized granule cells form an incipient dysgranular layer IV, and the transition of layers V and VI is gradual and almost imperceptible (Figure 2E). One neuroanatomist additionally included more posterior regions (referred to as Ph1-3) in their annotations of PHC (see Figure 4). These isocortical regions were characterized by a fully organized granular layer IV, though its prominence varied among the single subregions (see Supplementary Information for more details on regions Ph1-3).

All neuroanatomists placed PHC posterior to the ERC, lateral to the parasubiculum, and medial to the fundus of the posterior collateral sulcus extending as far as the posterior tip of the corpus callosum. Similar to the medial ERC border, the medial PHC border was annotated with high consistency in midsections (i.e., at the level of the hippocampal body; Figure 3). Figure 3 and Figure 4 additionally show the high disagreement in border placement in the posterior and lateral extent of PHC. Note that if the Ph subregion is not considered, which has been proposed to be part of the human PHC only recently (Stenger et al., 2022) and was annotated only by one neuroanatomist, the agreement among neuroanatomists on the PHC would be higher.

## Discussion

In this study, we give an overview of the cytoarchitectonic definitions of MTL cortex subregions (ERC, BA35 and 36, and PHC), aiming to examine and understand the underlying reasons for overlapping and diverging annotations made by four independent neuroanatomists. We find highest agreement of border placement for ERC and lowest agreement for PHC, while annotations of BA35 and BA36 show intermediate overlap. Importantly, the degree of agreement on definitions of cytoarchitectonic features does not necessarily correspond to agreement in border placement, since even structures with substantial agreement in described seminal features could show highly diverging annotations and vice versa. This partial mismatch of annotations and formal cytoarchitectonic definitions is due to additional factors influencing inter- and intra-rater reliability of neuroanatomical annotations as discussed below. Despite these differences, an extensive overlap of annotations among neuroanatomists is observed. Overall, the presented dataset of histological annotations uniquely informs *in vivo* neuroimaging research.

### Various factors influence the differences among neuroanatomists’ annotations

Based on the present results, we identify several factors which appear to influence the (dis-)agreement in annotations among neuroanatomists, underlining the complexity of annotating MTL cortex subregions.

First, disagreements are partially caused by varying definitions of features of MTL cortex subregions, as discussed above. These differences may partially stem from different traditions, backgrounds, and research foci of the neuroanatomists. For example, neuroanatomical schools may put particular emphasis on the subregions’ structural or functional connectivity, receptor architecture, or chemoarchitecture. Subsequently, this may lead to different perceptions and interpretations of histological information. For instance, we observed that while neuroanatomists agreed on the formal definition of different types of cortex, they did not allocate these to the same MTL cortex subregions. An example is proisocortex and the disagreement on whether this includes only BA35 or both BA35 and BA36. In several cases, the proisocortical BA35b of one and the proisocortical BA36 of another neuroanatomist were delineated almost identically. Additional variability may be introduced by the fact that neuroanatomical schools differ in their use of gross anatomical information for their annotations. This ranges from explicitly considering gross anatomy for border placement, using this information for orientation purposes, to dismissing gross anatomical landmarks and solely relying on histological information.

Second, we observed that annotations diverged in transitional areas between structures. Transition sharpness clearly differs within the MTL cortex, influencing the overlap of annotations. For example, the transition between ERC and hippocampal structures is relatively salient, leaving little room for uncertainty and leading to higher agreement among neuroanatomists. Meanwhile, there are transitional areas, such as that between BA35 and BA36, where the features of multiple subregions are expressed. As the neuroanatomists may weigh these features differently, they may come to different conclusions and place the respective border at different locations. This ambiguity does not only impact inter-rater reliability: neuroanatomists also highlighted that intra-rater reliability (i.e., one neuroanatomist rating the same specimen twice) would likely be lower in these transitional areas.

Third, the neuroanatomists’ certainty and, in turn, the agreement of annotations, is influenced by aspects related to the chosen sectioning procedure. The difficulty of annotating transitional zones may increase when borders between structures do not run strictly parallel or perpendicular to the provided sections. Even when using a consistent plane of sectioning (i.e., orthogonal to anterior commissure-posterior commissure (AC-PC) line) the irregular surface of the MTL – alongside the obliquity of sulci and other gross anatomical landmarks – can add complexity and ambiguity to the interpretation of boundaries. In such cases, it is possible that sections of one subregion are interrupted by portions of other types of cortices (e.g., appearance of olfactory cortex surrounded by ERC). In the present dataset, slices were cut perpendicular to the long axis of the hippocampus to aid the translation to thick-sliced *in vivo* MRI. For most neuroanatomists this is different from the common coronal sections cut perpendicular to the AC-PC line direction, which may have decreased certainty in border placement as well.

While we observed disagreement of annotations in transitional zones in all directions, transitions between sections (i.e., along the longitudinal axis) appear to cause more uncertainty than those within sections (i.e., on the medial to lateral axis). This may be because continuous information is only available within histological slices, making it more difficult to perceive differences between MTL cortical regions along the longitudinal axis. It is likely that this effect was exacerbated since three neuroanatomists were initially provided with histological slices spaced 5 mm apart, making subtle changes in cytoarchitectonic features more difficult to detect.

A fourth influencing factor that was pointed out by the neuroanatomists is the historical focus (or lack of) on some subregions over the others. The PHC, for example, is historically understudied in its seminal cytoarchitectonic features, connectivity, and borders with other structures. Most research of the PHC has been done in non-human primates (e.g., Blaizot et al., 2004; Bonin & Bailey, 1947; Suzuki & Amaral, 1994) and translational studies have only recently emerged (e.g., Stenger et al., 2022). Thus, less information is available on the cytoarchitectonic features of the human PHC (Ding et al., 2016; Ding & Van Hoesen, 2010; Mai et al., 2015; von Economo & Koskinas, 1925), resulting in lower agreement in parcellations.

Fifth, neuroanatomists pointed out that annotating at sulcal fundi is generally challenging as most cellular layers are compressed (see e.g., von Economo & Koskinas, 1929). This factor is relevant to the annotation of the border between BA35 and BA36 as well as the lateral border of PHC as both are often located in proximity to the fundus of the collateral sulcus.

Finally, neuroanatomists indicated that their uncertainties were amplified in the third specimen due to severe neurodegeneration of the MTL cortex. Cell loss that particularly affects certain neuronal populations and cortical layers distorts the overall cytoarchitectonic appearance of the cortex, potentially posing a significant challenge for neuroanatomical annotations. The uncertainty regarding the exact border locations may have contributed to higher disagreement across all annotated subregions in this specific specimen.

Performing annotations as presented in this study requires neuroanatomists to integrate various features of the provided specimen. Considering the aforementioned challenges leading to subjective uncertainty and disagreement among neuroanatomical schools, the present delineations of MTL cortex subregions exhibit impressively high overlap, particularly in the ERC and BA35.

### Implications for *in vivo* neuroimaging research

For human neuroimaging research, the precise and harmonized understanding of neuroanatomical entities ranging from small brain regions (such as MTL cortex subregions) to whole functional brain networks is crucial for synergistic work within and across research fields. The validity of labels applied in neuroimaging studies stems from a histologically-defined ground truth, and is key for integration and comparison of findings across a broader literature and for developing theories of brain structure, function, and pathology. An example for such collaborative processes is the investigation of the complex mechanisms underlying cognitive decline in neurodegenerative diseases such as Alzheimer’s disease. It requires the integration of insights from a wide range of research fields, ranging from *in vivo* functional MRI studies of functional specialization and/or gradients within the MTL, *in vivo* positron emission tomography studies to quantify pathology burden at different disease stages, to *ex vivo* structural MRI to explore the earliest neurodegenerative effects of Alzheimer’s disease pathology. Combining this multitude of research foci, modalities, and backgrounds requires a common language and harmonized neuroanatomical definitions. Thus, the presented overview of cytoarchitectonic features of MTL cortex subregions from four different neuroanatomical laboratories aids in further advancing human neuroimaging research in all of its applications and modalities.

The need for harmonization in neuroimaging is demonstrated in the variety of MRI segmentation protocols dedicated to the MTL (Berron et al., 2017; Feczko et al., 2009; Insausti et al., 1998; Kivisaari et al., 2020; Olsen et al., 2013; Pruessner et al., 2002; Wisse et al., 2017; Xie, Wisse, Pluta, Flores, et al., 2019; Yushkevich, Pluta, et al., 2015). Our study shows that while there is notable agreement among neuroanatomists in the cytoarchitectonic features referenced, there is currently no single neuroanatomical ground truth for the delineation of MTL cortex subregions on histological slices. As segmentation protocols for *in vivo* MRI commonly rely on a single neuroanatomical school, profound differences are inevitable. Consequently, *in vivo* neuroimaging studies of the MTL often lack comparability, negatively impacting their generalizability and usefulness for meta-analyses. To resolve this issue, harmonized definitions of MTL cortex subregions are needed for neuroimaging. Our present study sets an important foundation for this effort by providing an overview of how MTL cortex subregions are defined and delineated by different neuroanatomical schools. The Hippocampal Subfields Group is currently working towards establishing a harmonized segmentation protocol of the MTL cortex based on the presented findings.

### Strengths and limitations

A major strength of the present study is the integration of input from four leading expert neuroanatomists in the field, forming a first-of-its-kind annotated dataset of MTL cortex subregions in three histological specimens representing varying demographic backgrounds and anatomical variability. Yet, the limitations of the study must be considered.

First, no quantitative investigation of the differences in annotations was performed. Due to the manifold factors that influence the annotations of the neuroanatomists, we believe that it is not warranted to assess the differences in annotations quantitatively. Even more, focusing on quantitative measures of inter-rater agreement may lead to false conclusions and obscure the fine-grained picture that we are describing in this study. Instead, we show that considering the complexity of this work is of utmost importance when investigating the fine-grained neuroanatomy of the MTL cortex.

Second, the generalizability of our results is limited as the presented information is based on a small sample of three specimens. It has been reported that the location of MTL cortex subregions varies with collateral sulcus depth and ramification (Ding & Van Hoesen, 2010; Insausti et al., 1998). Although we included samples with different collateral sulcus depths and ramifications to account for this to some extent, a sample size of three specimens is clearly insufficient to fully characterize the role of anatomical variants in MTL subregion delineations. Unfortunately, because of the time and effort required (∼9 hours per specimen) to generate a dataset comprising the entirety of macroscopic MTL cortex phenotypes it was not possible to perform the current comparison between neuroanatomical schools in a larger sample. Nevertheless, we believe the characterization of the three specimens in this study still provide valuable insights.

Third, generalizability may also be impaired as we included “only” four neuroanatomists. There may be experts in the field who rely on neuroanatomical procedures that were not detailed here (e.g., intracellular filling). However, we again note that those techniques would be in principle limited to a small amount of the volume of the whole MTL cortex. Still, to our knowledge, the shown synthesis of neuroanatomical expertise goes beyond the efforts of other studies in the field, substantially improving upon the diverging opinions.

Finally, while Nissl-stained sections appear to provide optimal cytoarchitectural boundaries and appear to be reliable for performing mass parcellations (Williams et al., 2023), it has been suggested that Nissl-stained sections should be used in combination with immunochemical markers to annotate complex and transitional regions (e.g., Ding et al., 2009; Ding & Van Hoesen, 2010). Thus, the fact that we utilized Nissl-stained sections may have affected the certainty of some neuroanatomists negatively. However, the additional use of complementary techniques would not disambiguate the borders for all neuroanatomists. The time and effort required to implement such techniques, thus, do not justify their use.

## Conclusion

In summary, our detailed histological investigation shows that, aside from some disagreements, there is high overlap of definitions and delineations of subregions in the MTL cortex. These findings not only advance our neuroanatomical understanding of this important brain region but also provide a rich foundation for all neuroimaging research investigating the human MTL. Moreover, the in-depth understanding of disagreements among neuroanatomists likely translates to other brain regions, where similar degrees of disagreement are potentially present. This is evident for the hippocampus, where differences in annotations were observed among neuroanatomists (Olsen et al., 2019) and for neocortical regions as evidenced by different automated segmentations for *in vivo* MRI (Huizinga et al., 2021; Quilis-Sancho et al., 2020), potentially reflecting underlying differences in cytoarchitectonic definitions as well.

## Acknowledgements

We would like to acknowledge the donors and their family members for making this work possible. Special thanks go to the Marcus Wallenberg foundation for international scientific collaboration (2020.0004) and the Wenner-Gren foundation (ESh2020-002) for their generous support. This work was supported by NIH grants R01-AG070592, P01AG066597 and P30AG072979. Additionally, this study was supported by MultiPark - A Strategic Research Area at Lund University and by the Intramural UCLM grant GRIN 31097.

C.J.H. was supported by funding from the Biotechnology and Biological Sciences Research Council (BB/V010549/1). R.K.O. was supported by funding from the Canadian Institutes of Health Research (CIHR PJT162292) and the Alzheimer Society of Canada. T.T.T. was supported by the National Institute on Aging NRSA (F32AG071263), and the Alzheimer’s Association (AARFD-21-852597). P.Y. was supported by NIH grants P30 AG072979 and RF1 AG056014. O.K. and K.A. received funding from the European Union’s Horizon 2020 Research and Innovation Programme under the Specific Grant Agreement No. 945539 (Human Brain Project SGA3). J.C.A. was supported by NIH grants R01AG057672 and RF1AG072056. R.I. was supported by the Intramural UCLM grant GRIN 31097.

The funding sources had no role in the design and conduct of the study; in the collection, analysis, interpretation of the data; or in the preparation, review, or approval of the manuscript.

## Conflict of interest statement

The authors of this manuscript have nothing to declare.

## Data availability statement

Data can be shared upon completion of the harmonized protocol for the MTL cortex in a subsequent manuscript.

## Supplementary material

### Supplementary methods

The cloud-based digital histology archive was developed by the Penn Image Computing and Science Laboratory (PICSL), University of Pennsylvania, USA (https://picsl.upenn.edu/). The PICSL Histology Annotation Server (PHAS) allows the visualization and annotation of large serial histology datasets. Thus, a centralized data repository is easily produced. See https://github.com/pyushkevich/histoannot for more information. sFigure 1 shows an overview of the interface.

**sFigure 1.**
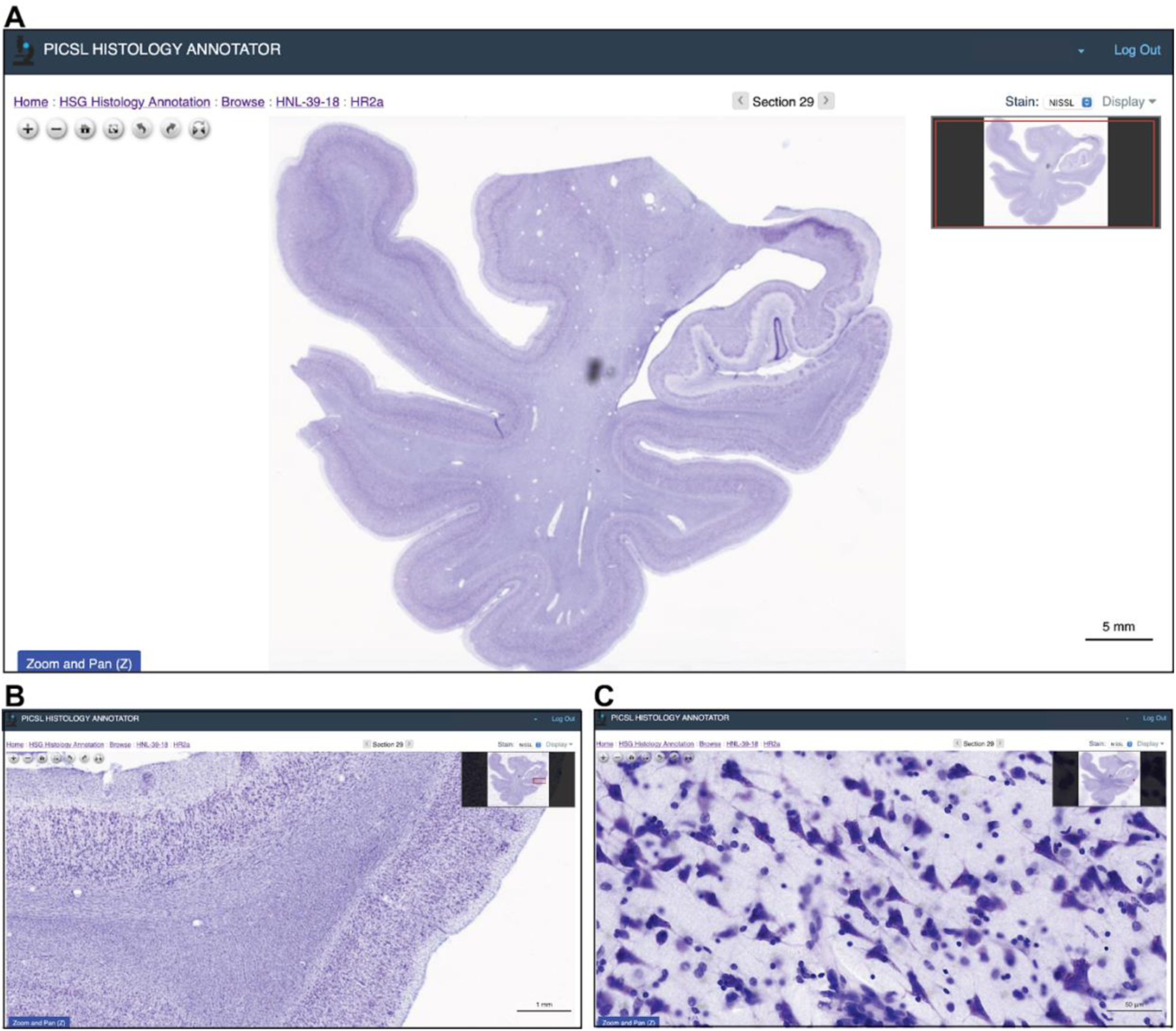
Interface of the PICSL Histology Annotation Server. A shows the overview page of a histology slice including scale bar and section specification while giving options to zoom and rotate images. B and C show the same slice displayed at higher zoom factors.

### Supplementary results

**sTable 1.** Differences in the nomenclature of the cortical layers of the ERC among neuroanatomists.

| R.I. (1995) | O.K. | J.C.A. | S.L.D. | Braak (1972) |
| --- | --- | --- | --- | --- |
| Layer I | Stratum<br>moleculare (1) | Layer I | Layer I | mol |
| Layer II | Stratum stellare<br>(2) | Layer II | Layer II | Pre-α |
| Layer III | Stratum<br>pyramidale (3) | Layer IIIa<br>Layer IIIb | Layer III | Pre-β<br>Pre-γ |
| Layer IV, Lamina<br>dissecans | Substratum<br>dissecans (3 diss) | Lamina dissecans<br>(when present) | Layer IV (Diss) | diss |
| Layer Va & Vb | Stratum<br>magnocellulare<br>(4) | Layer IV | Layer V | Pri-α |
| Layer Vc | Substratum<br>dissecans (4 diss) | Lamina dissecans<br>(when present) |  |  |
| Layer VI | Stratum<br>parvocellulare | Layer V | Layer VIa | Pri-β |
| | Stratum<br>multiforme | Layer VI | Layer VIb | Pri-γ |

Note that the terms or layer names in each column do not necessarily align with other columns (i.e., the number of layers is correct but names/representations may be different).

## Notes

### Competing Interest Statement

The authors have declared no competing interest.

### Summary of Updates

Updated author list

